# Activity-dependent post-translational regulation of palmitoylating and de-palmitoylating enzymes in the hippocampus

**DOI:** 10.1101/2022.09.12.507668

**Authors:** Danya Abazari, Angela R Wild, Tian Qiu, Bryan C. Dickinson, Shernaz X Bamji

## Abstract

Activity-induced changes in protein palmitoylation can regulate the plasticity of synaptic connections, critically impacting learning and memory. Palmitoylation is a reversible post-translational modification regulated by both palmitoyl-acyl transferases that mediate palmitoylation and palmitoyl thioesterases that depalmitoylate proteins. However, it is not clear how fluctuations in synaptic activity can mediate the dynamic palmitoylation of neuronal proteins. Using primary hippocampal cultures, we demonstrate that synaptic activity does not impact the transcription of palmitoylating / depalmitoylating enzymes, changes in thioesterase activity, nor post-translational modification of the depalmitoylating enzymes, ABHD17 and APT2. In contrast, synaptic activity does mediate post-translational modification of the palmitoylating enzymes ZDHHC2, ZDHHC5, and ZDHHC9 (but not ZDHHC8) to influence protein-protein interaction, enzyme stability and enzyme function. Post-translational modifications of ZDHHC enzymes were also observed in the hippocampus following fear conditioning. Together, our findings demonstrate that signaling events activated by synaptic activity largely impact activity of the ZDHHC family of palmitoyl-acyl transferases with less influence on the activity of palmitoyl thioesterases.

## Introduction

The formation and remodeling of synaptic contacts requires the precise distribution and trafficking of proteins to specialized compartments. While post-translational phosphorylation of synaptic proteins has been well-studied and shown to play a key role in regulating synaptic plasticity (Anggono and Huganir, 2012; Esteban et al., 2003; Hayashi, 2021; Lee, 2006; Lua et al., 2010; Man et al., 2003; Woolfrey and Dell’Acqua, 2015), more recent studies have demonstrated that other post-translational modifications, including protein *S*-acylation, can be equally important for the strengthening and weakening of synaptic connections.

The most prominent form of *S*-acylation, and the most common post-translational lipid modification in the brain, is *S*-palmitoylation (hereafter called palmitoylation). Palmitoylation involves the reversible addition of palmitoyl moieties to selected cysteine residues via thioester bonds, increasing protein hydrophobicity and the affinity for plasma membranes. This reaction is catalyzed by a family of 23 palmitoylating ZDHHC enzymes, and reversed by a subset of proteins in the serine hydrolase superfamily (Main and Fuller, 2022). Approximately 41% of all known synaptic proteins can be palmitoylated (Sanders et al., 2015) including ion channels (Pei et al., 2018), SNARE proteins (Greaves et al., 2010; He and Linder, 2009), scaffold proteins (El-Husseini et al., 2000; Purkey et al., 2018; Topinka and Bredt, 1998; Vallejo et al., 2017), signaling molecules (Rocks et al., 2005), and neurotransmitter receptors including AMPA, NMDA and GABA receptor subunits (Hayashi et al., 2009, 2005; Lin et al., 2009; Resh, 2006; Thomas and Huganir, 2013).

Notably, several studies have shown that synaptic proteins can be differentially palmitoylated in response to synaptic activity and that the dynamic palmitoylation of synaptic proteins is essential for synapse plasticity. Using an unbiased proteomic approach, our lab has identified 121 proteins (56 synaptic proteins) that are differentially palmitoylated in response to fear conditioning, and that a subset of these are also differentially palmitoylated in response to increased synaptic activity in primary hippocampal cultures (Nasseri et al., 2021). Moreover, work from our lab and others have shown that increased synaptic activity can increase the palmitoylation of PSD-95 (Noritake et al., 2009), AKAP79/150 (Keith et al., 2012; Woolfrey et al., 2015), δ-catenin (Brigidi et al., 2015, 2014), and PRG-1/LPRR4 (plasticity-related gene 1/ lipid phosphate phosphatase-related protein type 4) (Nasseri et al., 2021), and that the palmitoylation of these proteins are essential for the recruitment and retention of AMPARs to the synaptic membrane and the strengthening of synaptic connections. While these studies demonstrate that dynamic protein palmitoylation is important for synaptic plasticity, it is unclear *how* changes in synaptic activity can alter protein palmitoylation.

According to BrainPalmSeq, an RNAseq database tool developed in our lab, the majority of ZDHHC enzymes are expressed in hippocampal excitatory neurons (Wild et al., 2021). While many ZDHHCs reside within the somatic Golgi where they constitutively palmitoylate proteins, several are localized to dendrites where they can locally and dynamically palmitoylate synaptic proteins. These include ZDHHC2 (Fukata et al., 2013), ZDHHC5 (Brigidi et al., 2015; Thomas et al., 2012), ZDHHC8 (Thomas et al., 2012), ZDHHC9 (Shimell et al., 2019), ZDHHC14 (Sanders et al., 2020) and ZDHHC15 (Shah et al., 2019). Although identification of the full family of depalmitoylating enzymes is still underway, a subset are known to be expressed in neurons (Wild et al., 2021) and localized to neuronal processes. These include APT1 and APT2 (Milde and Coleman, 2014; Shen et al., 2022), PPT1 which is targeted to axons (Ahtiainen et al., 2003; Kim et al., 2008) and the more recently discovered α/β-hydrolase domain-containing protein 17 members (ABHD17A, 17B, and 17C), which have numerous post-synaptic substrates, including PSD-95, BK channels and NRAS (Lin and Conibear, 2015; McClafferty et al., 2020; Yokoi et al., 2016). These palmitoylating and depalmitoylating enzymes are therefore well positioned to mediate dynamic substrate palmitoylation that occurs following changes in synaptic activity.

While increasing evidence suggests that synaptic activity leads to the differential palmitoylation of neuronal proteins, and that palmitoylation is important for the strengthening and weakening of synaptic connections, the mechanisms by which this occurs are largely unknown. In this study we demonstrate that activity-induced changes in protein palmitoylation are largely driven by the dynamic function of palmitoylating enzymes as opposed to depalmitoylating enzymes. Increasing synaptic activity in primary hippocampal cultures mediates the post-translational modification of palmitoylating enzymes, ZDHHC2, ZDHHC5 and ZDHHC9, but not of ZDHHC8 or the depalmitoylating enzymes, APT1 and ABHD17. We further demonstrate that these post-translational modifications are essential for ZDHHC enzyme stability, protein interactions, as well as enzymatic activity. Notably, similar changes in posttranslational modifications of these ZDHHC enzymes occurred 1 hour after fear conditioning, highlighting the importance of ZDHHC modifications in regulating synapse function *in vivo*. Together, these data suggest that the differential palmitoylation of synaptic proteins upon synaptic stimulation is mediated by through post-translational modifications of ZDHHC enzymes that in turn regulate enzyme stability and function.

## Results

### ZDHHC enzyme transcription is largely unchanged after increased synaptic activity

Strong evidence from multiple studies has shown that changes in synaptic activity can dramatically alter the transcriptional profiles of neuronal proteins (Flavell and Greenberg, 2008; West et al., 2002). Notably, protocols that induce long-term potentiation (LTP) can also alter transcription of numerous neuronal genes (Bliim et al., 2019; Tyssowski et al., 2018). To determine whether synaptic activity alters the transcription profile of ZDHHC enzymes as a means to regulate dynamic substrate palmitoylation, we increased synaptic activity in cultured hippocampal neurons using a well-established chemical LTP (cLTP) protocol involving a brief 3 min incubation with 200 μM glycine in the absence of Mg^2+^ (Lu et al., 2001), and quantified mRNA transcripts for the 23 ZDHHC enzymes (Uniprot reviewed) 40min, 2h and 24h later. While cLTP did not significantly alter transcription of the majority of the ZDHHCs at each time point (Figure 1), expression of *Zdhhc2, Zdhhc8*, and *Zdhhc22* was significantly reduced, and expression of *Zdhhc11* increased 24 hours following cLTP induction. As activity-induced changes in protein palmitoylation have been shown to occur at much earlier timepoints (Brigidi et al., 2015; Nasseri et al., 2021; Noritake et al., 2009), it is unlikely that changes in the transcription of these few ZDHHC genes are grossly responsible for alterations in protein palmitoylation. We therefore next investigated alternative regulatory mechanisms that might alter ZDHHC function minutes to hours after synaptic stimulation, when activity-induced changes in substrate palmitoylation are known to occur (Nasseri et al., 2021).

**Figure 1.**
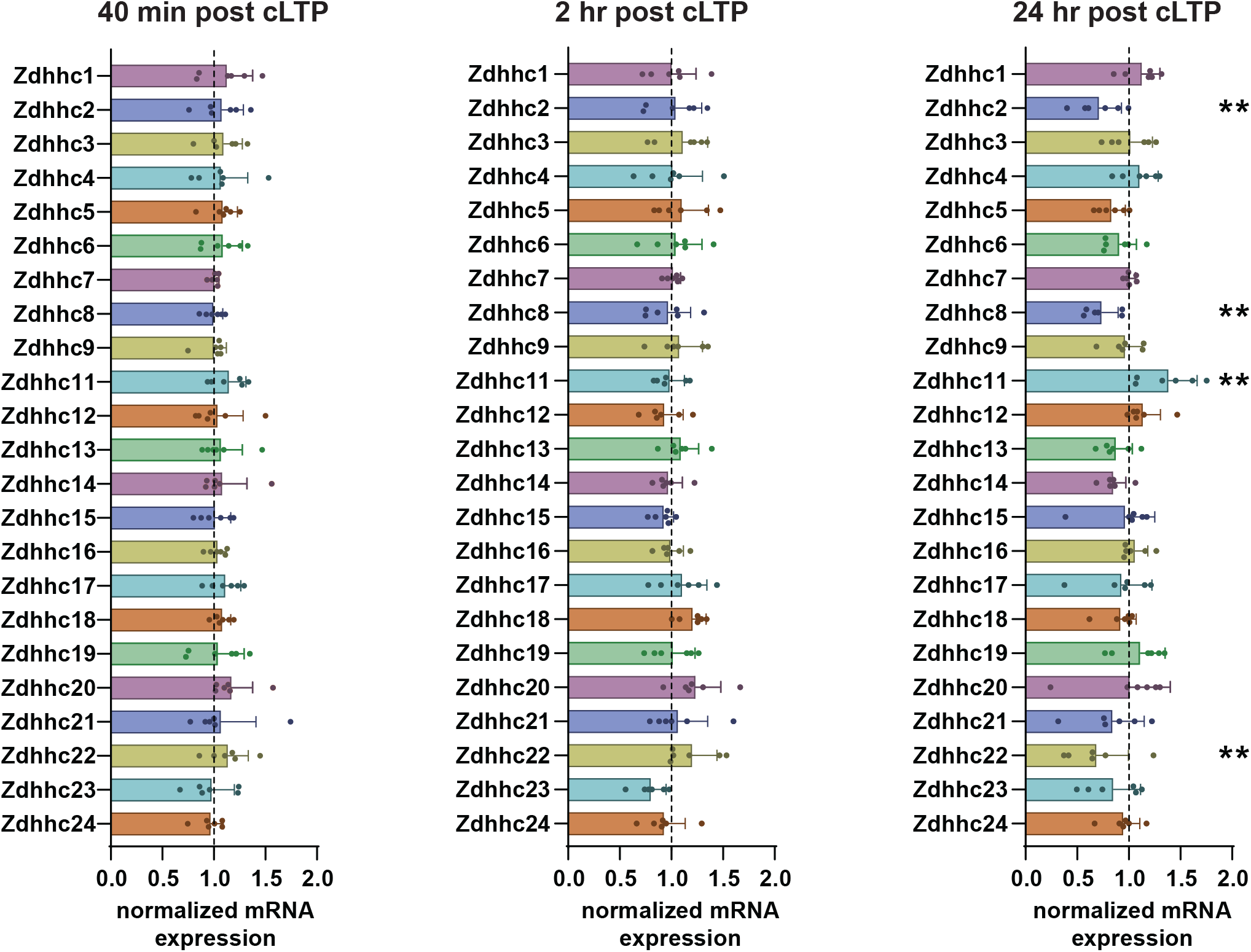
ZDHHC mRNA levels are not changed 40 min and 2 hrs post cLTP. qRT-PCR of the 23 ZDHHC enzymes from primary hippocampal cultures 40min, 2 hrs or 24 hrs following cLTP. Values are normalized to mock-cLTP treated control. **P<0.01 (unpaired student’s *t*-test vs mock-cLTP control). Results are mean ± s.e.m. with individual data points shown. N = 6 independent hippocampal cultures per condition.

### ZDHHC antibody validation for biochemical study of endogenous ZDHHCs

To assay activity-induced changes in endogenous ZDHHCs, we first needed to identify commercially available antibodies that detected the ZDHHC with high specificity. Upon testing available antibodies for all 23 ZDHHCs enzymes, good antibodies were identified for ZDHHC2, ZDHHC5, ZDHHC6, ZDHHC8 and ZDHHC9 (Supplementary Fig. 1). We chose to further study ZDHHC2 and ZDHHC5 as they have been shown to regulate activity-induced palmitoylation of synaptic proteins (Brigidi et al., 2015, 2014; Fukata et al., 2013; Noritake et al., 2009), as well as ZDHHC8 and ZDHHC9 whose function is disrupted in a subset of patients with schizophrenia (Mukai et al., 2004) and X-linked intellectual disability (Baker et al., 2015; Raymond et al., 2007), respectively. Notably, these four enzymes are highly expressed in hippocampal neurons (Wild et al., 2022), have well defined roles in regulating synaptic function and localize to neuronal dendrites and spines where they are appropriately positioned to mediate rapid, dynamic changes in the palmitoylation of synaptic proteins in response to changes in synaptic activity (Brigidi et al., 2015; Fukata et al., 2013; Shimell et al., 2019, 2021; Thomas et al., 2012, 2013; Woolfrey and Dell’Acqua, 2015).

### Activity-dependent ZDHHC5 degradation is regulated by phosphorylation of a Polo box motif

Our lab has previously shown that synaptic activity increases palmitoylation of the cadherin binding protein, δ-catenin (CTNND1), and that this is mediated by the dephosphorylation of ZDHHC5 on tyrosine residue 533 and the subsequent internalization of ZDHHC5 from the plasma membrane (Brigidi et al., 2015). To get a more fulsome understanding of how synaptic activity can impact ZDHHC5 function, we monitored cLTP-induced changes in protein turnover and post-translational modifications. We focused on the post-translational modifications phosphorylation and palmitoylation, which are both highly dynamic and have considerable influence over synaptic protein function and localization (Ji and Skup, 2021; Lee, 2006). Furthermore, kinases and phosphatases that mediate phosphorylation are highly responsive to synaptic activity (Woolfrey and Dell’Acqua, 2015), while certain ZDHHCs are known to engage in palmitoylation cascades whereby ZDHHC enzymes are themselves palmitoylation substrates for other ZDHHC enzymes that control their function (Abrami et al., 2017; Plain et al., 2020). We observed a substantial (> 50 %) reduction in the total protein levels of ZDHHC5 40 mins post-cLTP, which recovered slightly at 2 hours but did not return to baseline after 24 hrs (Figure 2A). We also observed an activity-dependent increase in palmitoylation of ZDHHC5 using an acyl-RAC assay (Badrilla, UK) at 40 mins and 24 hrs post cLTP, when the palmitoylated fraction was normalized to total ZDHHC5 (Figure 2B). Additionally, when using a phospho-protein affinity enrichment assay (PhosphoProtein Purification Kit; Qiagen) to assess the overall change in the amount of phosphorylated ZDHHC5, we observed a significant overall increase in the phosphorylated fraction of ZDHHC5 (when normalized to total protein input; Figure. 2C). As this increase in ZDHHC5 phosphorylation initially appears to be counter to our previous observations of decreased Tyr phosphorylation following cLTP (Brigidi et al., 2015), we further investigated which phospho-residues may be responsible for the overall net increase in ZDHHC5 phosphorylation, and how this might be related to the substantial decrease in total ZDHHC5 protein.

**Figure 2.**
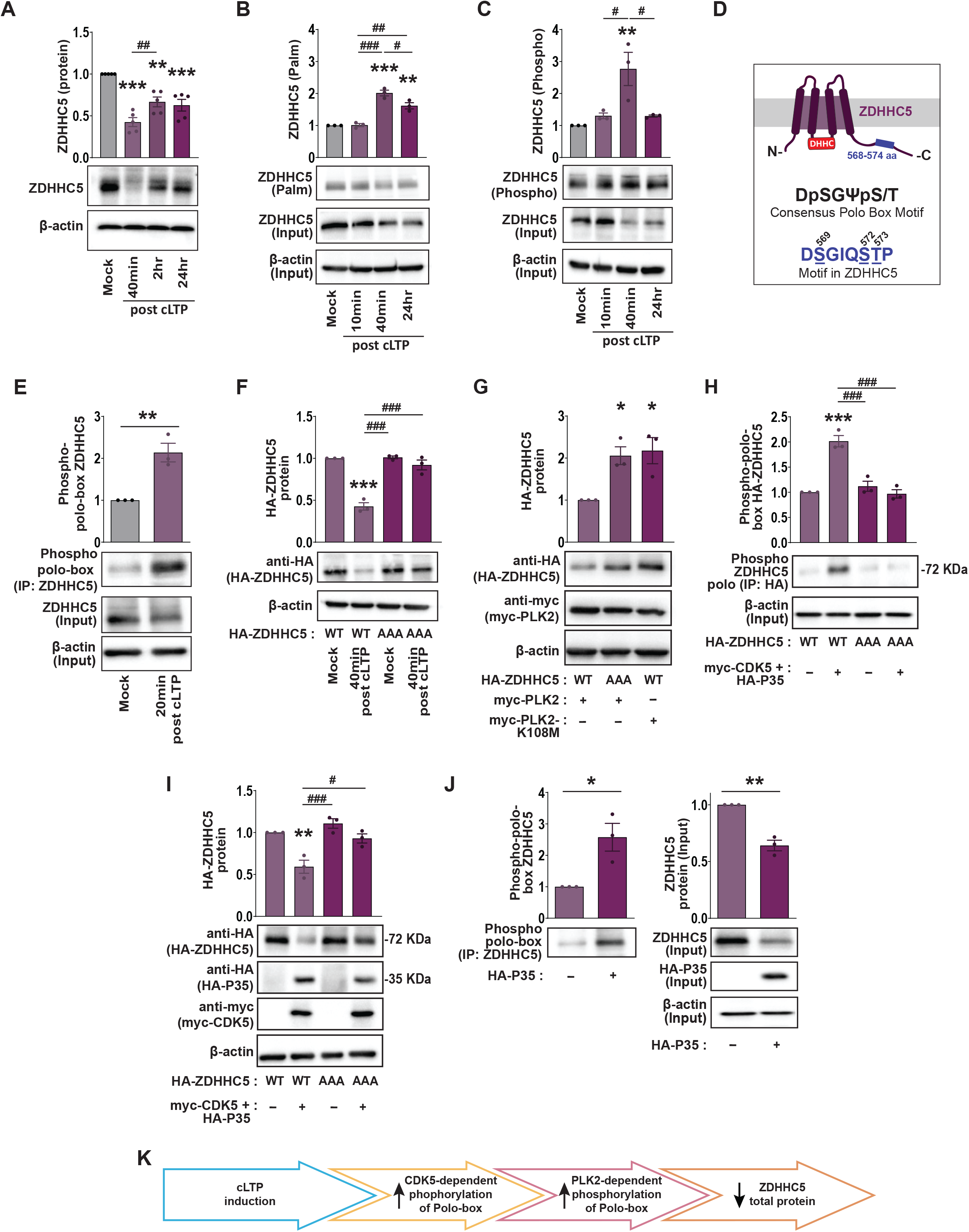
cLTP-induced phosphorylation of ZDHHC5 polo-box domain impacts ZDHHC5 stability. **(A)** Western blot analysis of ZDHHC5 protein levels in primary hippocampal neuron cultures 40 min, 2 hrs and 24 hrs following cLTP. **(B)** Acyl-Rac assay showing palmitoylated ZDHHC5 and overall ZDHHC5 protein (input) levels following cLTP. Palmitoylated ZDHHC5 values in graph derived from ZDHHC5 ‘palm’ normalized to ZDHHC5 ‘input’. **(C)** Lysates were run through the phospho-protein purification assay and Western Blots probed with ZDHHC5 antibody showing phosphorylated ZDHHC5 and overall ZDHHC5 protein (input) levels following cLTP. Phosphorylated ZDHHC5 values in graph derived from ZDHHC5 ‘phospho’ normalized to ZDHHC5 ‘input’. **(D)** Schematic of the sequence and position of polo-box motif within ZDHHC5 C-terminal tail. **(E)** Hippocampal culture lysates immunoprecipitated (IP) with a ZDHHC5 antibody and Western blots probed with a phopho-polo-box specific antibody. *P<0.05, **P < 0.01, ***P < 0.001 (unpaired student’s *t*-test). **(F)** Cells were nucleofected with ZDHHC5 shRNA to knock down ZDHHC5 together with HA-ZDHHC5 WT or HA-DHHC5 AAA mutant at time of plating. At 13-15 DIV, cultures were stimulated using cLTP treatment and HA-ZDHHC5 protein levels determined using Western blot analysis. **(G)** Cells were nucleofected with myc-PLK2 WT or kinase dead PLK2 (myc-PLK2-K108M) together with HA-ZDHHC5-WT or HA-ZDHHC5-AAA. ZDHHC5 or PLK2 protein levels were determined at 14 DIV by Western blotting. **(H)** Cells were nucleofected with HA-ZDHHC5-WT or HA-ZDHHC5-AAA plus myc-CDK5 and HA-P35. At 13-15 DIV HA-tagged proteins were immunoprecipitated (IP) with an HA antibody and Western blots probed with a phopho-polo-box antibody determine ZDHHC5 polo-box phosphorylation. **(I)** Cells were nucleofected with HA-ZDHHC5-WT or HA-ZDHHC5-AAA plus myc-CDK5 and HA-P35. At 14 DIV Western blot was performed to assay protein levels of all transfected constructs. *P<0.05, **P < 0.01, ***P < 0.001 vs first condition in bar chart. #P < 0.05, ##P < 0.01, ###P < 0.001 pairwise comparison as indicated (one-way ANOVA with Tukey’s post hoc test). **(J)** Cells were nucleofected with HA-P35. At 14 DIV ZDHHC5 was immunoprecipitated (IP) with an endogenous ZDHHC5 antibody and Western blots probed with a phopho-polo-box antibody to assay ZDHHC5 polo-box phosphorylation (left). The input from this assay was run on a Western blot and probed for endogenous ZDHHC5 and HA-P35 (right). *P<0.05, **P < 0.01, ***P < 0.001 (unpaired student’s *t*-test). Results are mean ± s.e.m. with individual data points shown. N = 3 - 5 independent hippocampal cultures per experiment. **(K)** Schematic of activity dependent changes in ZDHHC5 C-terminal polo-box regulation of ZDHHC5 total protein.

Using bioinformatics analysis of the C-terminal of ZDHHC5 we identified a sequence (DSGIQSTP) very similar to the consensus ‘Polo-box’ motif DpSGΨXpS/T (Ψ=hydrophobic residue, X= any residue, pS or pS/T= phosphoserine or threonine), so named due to the sequence being recognized by the ‘Polo domain’ present in polo-like kinases (Nakojima et al., 2003; Figure 2D). When dually phosphorylated on serine/threonine residues, this motif targets proteins for rapid ubiquitination and degradation (Ang et al., 2008; Arai et al., 2008; Moshe et al., 2004; Pak and Sheng, 2003; Seeburg et al., 2008). We therefore investigated whether cLTP increases ZDHHC5 phosphorylation on these serine/threonine residues and whether this regulates ZDHHC5 stability. Hippocampal culture lysates were immunoprecipitated with ZDHHC5 and blots probed with an antibody that specifically recognizes the phosphorylated Polo box motif (Baehr et al., 2016; Wang et al., 2018) 20 minutes following cLTP treatment. We observed a dramatic, 2-fold increase in the phosphorylation of Ser/thr residues in this motif despite a significant decrease in ZDHHC5 protein levels (input) in neurons treated with cLTP (Fig. 2E). To further ascertain whether the phosphorylation of this motif is required for activity-induced degradation of ZDHHC5, hippocampal neurons were transfected with ZDHHC5 shRNA to knockdown endogenous ZDHHC5, plus HA tagged either wildtype ZDHHC5 (HA-ZDHHC5-WT) or phospho-dead ZDHHC5 (HA-ZDHHC5-AAA), where Ser569, Ser572 and Thr574 in the Polo box motif were changed to alanines. While cLTP significantly decreased the expression of HA-ZDHHC5-WT, phospho-dead HA-ZDHHC5-AAA levels were unchanged (Fig. 2F), demonstrating that phosphorylation of this motif is required for degradation of ZDHHC5 following cLTP.

We next set out to determine how synaptic activity can regulate ZDHHC5 phosphorylation and subsequently the destabilization of ZDHHC5. Previous studies have shown that Polo-Like Kinase2 (PLK2) can phosphorylate residues within Polo-box motifs (Ang et al., 2008; Lee et al., 2011). We therefore determined whether PLK2 is involved in phosphorylation-dependent degradation of ZDHHC5. While overexpression of wild-type myc-tagged (myc-PLK2-WT) resulted in a decrease in HA-ZDHHC5-WT protein levels, myc-PLK2 had no effect on HA-ZDHHC5-AAA total protein (Fig. 2G). Moreover, the PLK2 kinase-dead mutant in which Lys 108 is mutated to a Met (myc-PLK2-K108M) mutant did not impact HA-ZDHHC5-WT protein levels (Fig. 2G), demonstrating that PLK2 mediates ZDHHC5 degradation through phosphorylation of the Polo box motif.

Prior to phosphorylation by PLK2, DpSGΨXpS/T containing peptides have shown to be phosphorylated by proline-directed kinases such as cyclin dependent kinases (CDKs; Hamanaka et al., 1995; Martin and Strebhardt, 2006; Seeburg et al., 2008; Thomas et al., 2016). Indeed, it is thought that CDK-mediated phosphorylation can prime proteins to be phosphorylated by PLK2 (Elia et al., 2003). Previous work has identified CDK5 as the priming kinase that phosphorylates STP motifs in the substrate protein, SPAR (Seeburg et al., 2008). To see whether CDK5 is involved in the Polo box phosphorylation and destabilization of ZDHHC5, hippocampal neurons were transfected with HA-ZDHHC5-WT or HA-ZDHHC5-AAA together with CDK5 and its neuronal-specific activator, P35 (Chae et al., 1997). Overexpression of myc-CDK5 and HA-P35 increased the phosphorylation of the Polo box domain (Fig. 2H) and decreased overall levels of HA-ZDHHC5-WT but not HA-ZDHHC5-AAA (Fig 2I). Finally, overexpression of HA-P35 alone was sufficient to activate endogenous CDK5 and increase phosphorylation of the Polo box motif within immunoprecipitated endogenous ZDHHC5 (Fig. 2J). This was accompanied by a decrease in total endogenous ZDHHC5 protein in the input fraction (Fig. 2J, right). We have therefore identified a mechanism by which PLK2, CDK5 and P35 regulate the stability of ZDHHC5 in neurons following synaptic activity (Fig. 2K). In line with our previous work, these results reveal that ZDHHC5 is highly responsive to synaptic activity and as such well positioned to mediate dynamic changes in palmitoylation of synaptic proteins.

### ZDHHC8 phosphorylation, palmitoylation and protein turnover are not affected by cLTP

We next investigated the effects of cLTP on the post translation regulation of ZDHHC8, which has a role in synaptic development (Mukai et al., 2008) and was found to be disrupted in a subset of patients with schizophrenia (Mukai et al., 2004). ZDHHC8 localizes to dendritic projections and is the closest homologue of ZDHHC5, with which it shares 60% sequence similarity and 50% identity. Furthermore, ZDHHC8 contains many signaling motifs in common with ZDHHC5, including a C-terminal PDZ binding domain (EISV; Thomas et al., 2012), tyrosine endocytic motif (YDNL; Brigidi et al., 2015), along with a Polo box like domain (DSGVYDT). We therefore determined if ZDHHC8 PTMs and protein turnover were altered by synaptic stimulation with cLTP. Surprisingly, no changes were observed in endogenous ZDHHC8 total protein levels 40 minutes, 2 hours, and 24 hours after cLTP (Fig. 3A), indicating that the reduction in ZDHHC8 mRNA we observed 24 hours post cLTP (Fig. 1) did not significantly alter total protein turnover. We next investigated if ZDHHC8 palmitoylation is modified by cLTP, as Cys residues 236 and 245 of the murine ZDHHC8 C-terminal have been shown to be palmitoylated (Collins et al., 2017). Accordingly, we detected palmitoylated ZDHHC8 in the palmitoylated fraction using the acyl-Rac assay, but this was not altered following cLTP (Fig. 3B). Finally, we did not observe changes in ZDHHC8 phosphorylation following cLTP treatment (Fig. 3C). Together, these results indicate that unlike ZDHHC5, ZDHHC8 may be less responsive to increased synaptic activity stimulated by cLTP, despite its localization to neuronal dendrites and synapses (Thomas et al., 2012).

**Figure 3.**
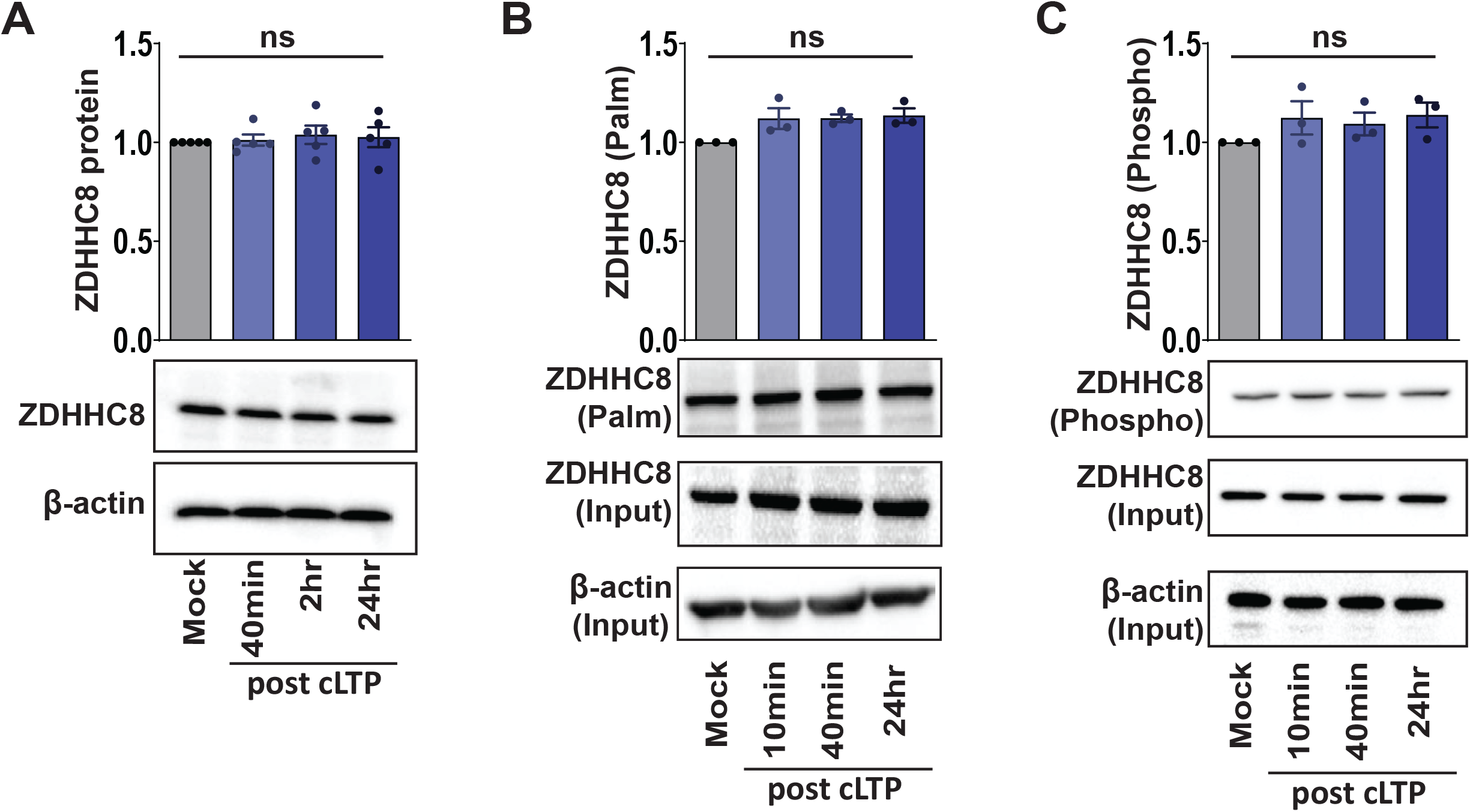
ZDHHC8 protein levels, palmitoylation and phosphorylation are unchanged following cLTP. **(A)** Western blot analysis of ZDHHC8 protein levels in primary hippocampal neuron cultures 40 min, 2 hrs and 24 hrs following cLTP. **(B)** Acyl-Rac assay showing palmitoylated ZDHHC8 and overall ZDHHC8 protein (input) levels following cLTP. Palmitoylated ZDHHC8 values in graph derived from ZDHHC8 ‘palm’ normalized to ZDHHC8 ‘input’. **(C)** Lysates were run through the phospho-protein purification assay and Western Blots probed with ZDHHC8 antibody showing phosphorylated ZDHHC8 and overall ZDHHC8 protein (input) levels following cLTP. Phosphorylated ZDHHC8 values in graph derived from ZDHHC8 ‘phospho’ normalized to ZDHHC8 ‘input’. ns = not significant (one-way ANOVA with Tukey’s post hoc test). Results are mean ± s.e.m. with individual data points shown. N = 3-5 independent hippocampal cultures per experiment.

### ZDHHC9 palmitoylation and substrate palmitoylation are decreased by cLTP

We recently demonstrated that disrupting ZDHHC9 function *in vitro* decreases both dendritic outgrowth and the formation of inhibitory synapses (Shimell et al., 2019). As above, we assayed the effects of cLTP on ZDHHC9 palmitoylation, phosphorylation and turnover. We found that while ZDHHC9 protein levels (Fig. 4A) and phosphorylation (Fig. 4C) were unchanged, palmitoylation of ZDHHC9 significantly decreased to 50% of baseline levels 10 min after cLTP and was maintained up to 24 hrs after cLTP treatment (Fig. 4B). Previous work has shown that ZDHHC enzymes are first palmitoylated on the cysteine residue in the DHHC domain before transferring palmitate (palmitic acid) to its substrate (Stix et al., 2020). To determine whether cLTP specifically decreases ZDHHC9 palmitoylation at this site, cells were transfected with either WT or a DHH<u>S</u>9 mutant (Cys 169 mutated to Ser). As expected, there was a decrease in the palmitoylation of WT-ZDHHC9 1 hr after cLTP (Fig 4D). Basal palmitoylation of DHH<u>S</u>9 was significantly reduced compared to WT-ZDHHC9 demonstrating that palmitoylation of this residue does indeed contribute to overall palmitoylation of ZDHHC9. Notably, ZDHHS9 palmitoylation did not decrease further following cLTP treatment, indicating that this residue is subject to activity-dependent depalmitoylation (Fig 4D).

**Figure 4.**
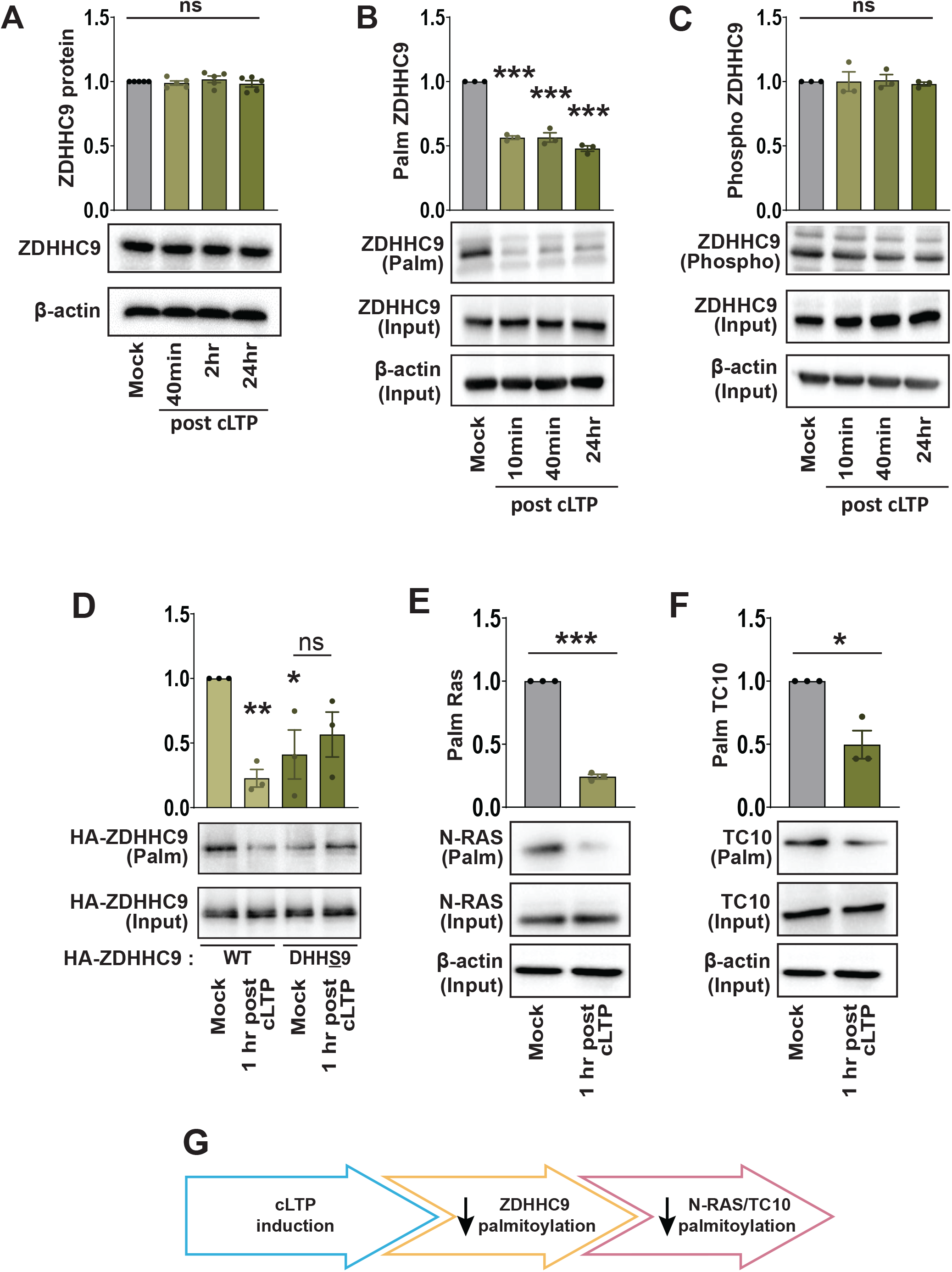
The palmitoylation of ZDHHC9 and its substrates are reduced following cLTP. **(A)** Western blot analysis of ZDHHC9 protein levels in primary hippocampal neuron cultures 40 min, 2 hrs and 24 hrs following cLTP. **(B)** Acyl-Rac assay showing palmitoylated ZDHHC9 and overall ZDHHC9 protein (input) levels following cLTP. Palmitoylated ZDHHC9 values in graph derived from ZDHHC9 ‘palm’ normalized to ZDHHC9 ‘input’. **(C)** Lysates were run through the phospho-protein purification assay and Western Blots probed with ZDHHC9 antibody showing phosphorylated ZDHHC9 and overall ZDHHC9 protein (input) levels following cLTP. Phosphorylated ZDHHC9 values in graph derived from ZDHHC9 ‘phospho’ normalized to ZDHHC9 ‘input’. **(D)** Cells were nucleofected at time of plating with ZDHHC9 shRNA to knock down ZDHHC9 together with HA-ZDHHC9-WT or HA-ZDHH<u>S</u>9. At 13-15 DIV, cultures were stimulated using cLTP treatment and HA-ZDHHC9 palmitoylation was determined using the Acyl-Rac assay. *P<0.05, ***P < 0.001 vs first condition in bar chart (one-way ANOVA with Tukey’s post hoc test). **(E, F)** At 14 DIV, cultures were stimulated using cLTP treatment and N-RAS **(E)** or TC10 **(F)** palmitoylation was determined using the Acyl-Rac assay. **(G)** Schematic showing activity-induced decrease in ZDHHC9 and substrate palmitoylation. ns = not significant. *P<0.05, **P < 0.01, ***P < 0.001 (unpaired student’s *t*-test). Results are mean ± s.e.m. with individual data points shown. N = 3-5 independent hippocampal cultures per experiment.

ZDHHC9 has several neuronal substrates including NRAS and TC10 (Shimell et al., 2019). To determine whether activity-induced depalmitoylation of ZDHHC9 impacts the palmitoylation of NRAS and TC10, we assayed their palmitoylation 1 hr after cLTP induction. There was a substantial decrease in the palmitoylation of these two proteins (Fig. 4E, F), suggesting that synaptic activity can decrease ZDHHC9 enzymatic activity by reducing its palmitoylation within the catalytic domain (Fig 4G).

### ZDHHC2 phosphorylation and PSD-95 interaction decrease after cLTP

ZDHHC2 localizes to hippocampal dendrites, where it cycles between recycling endosomes and the plasma membrane (Fukata et al., 2013). ZDHHC2 regulates the palmitoylation of the synaptic scaffold protein, PSD-95 (Fukata et al., 2013; Noritake et al., 2009), and AKAP79/150 (A-kinase anchoring protein; Woolfrey et al., 2015), thereby regulating the synaptic localization of AMPAR subunits and synapse strength. Similar to other ZDHHCs above, we interrogated activity-induced changes in ZDHHC2 protein turnover and post-translational modifications following cLTP treatment. While total ZDHHC2 protein levels (Fig. 5A), and ZDHHC2 palmitoylation (Fig. 5B) were unchanged, we observed a significant reduction in the phosphorylation of ZDHHC2 40 min and 24 hrs post cLTP (Fig. 5C).

**Figure 5.**
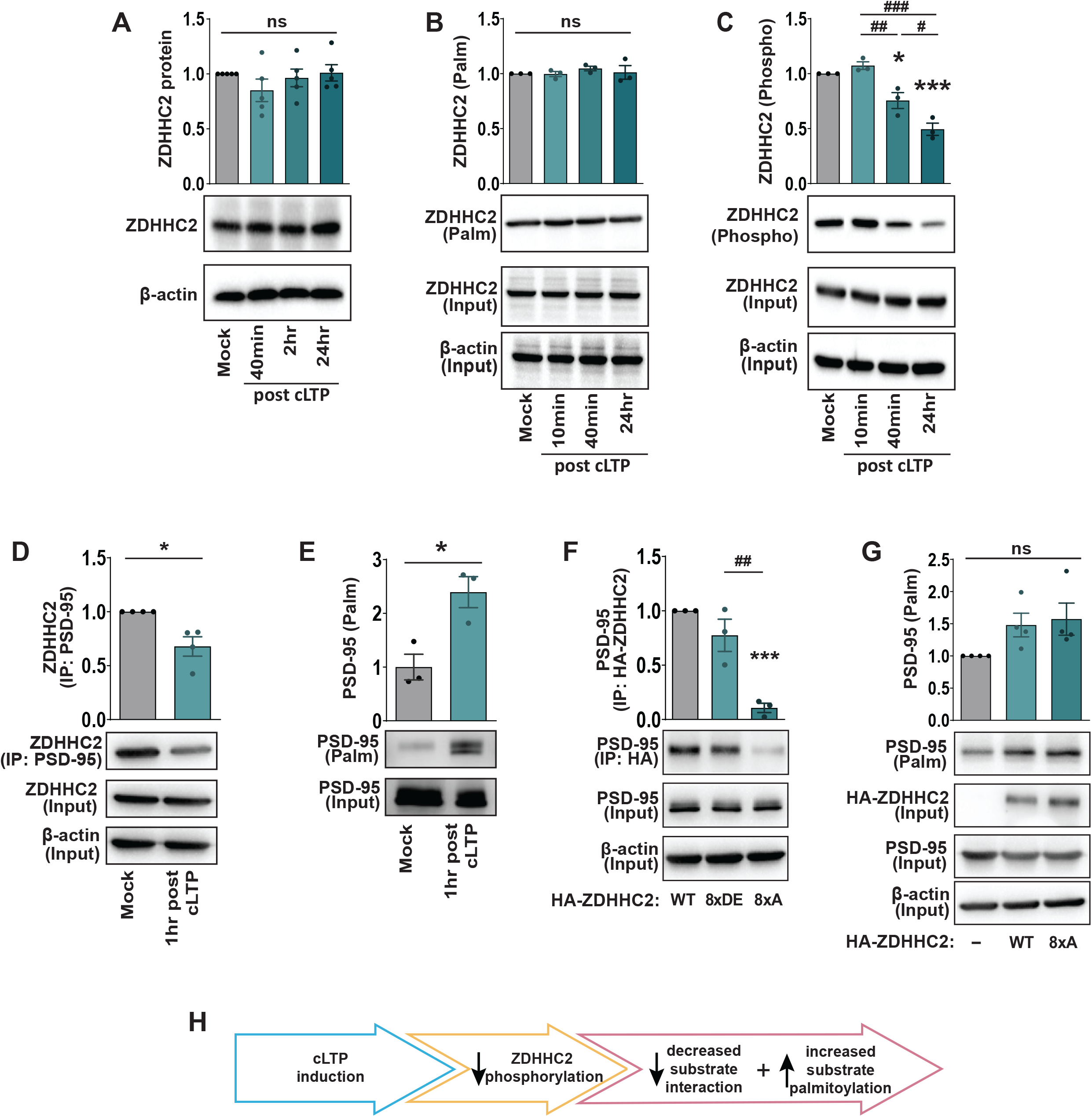
Decreased phosphorylation of ZDHHC2 following cLTP. **(A)** Western blot analysis of ZDHHC2 protein levels in primary hippocampal neuron cultures 40 min, 2 hrs and 24 hrs following cLTP. **(B)** Acyl-Rac assay showing palmitoylated ZDHHC2 and overall ZDHHC2 protein (input) levels following cLTP. Palmitoylated ZDHHC2 values in graph derived from ZDHHC2 ‘palm’ normalized to ZDHHC2 ‘input’. **(C)** Lysates were run through the phospho-protein purification assay and Western Blots probed with ZDHHC2 antibody showing phosphorylated ZDHHC2 and overall ZDHHC2 protein (input) levels following cLTP. Phosphorylated ZDHHC2 values in graph derived from ZDHHC2 ‘phospho’ normalized to ZDHHC2 ‘input’. ns = not significant. *P<0.05, ***P < 0.001 vs first condition in bar chart. #P < 0.05, ##P < 0.01, ###P < 0.001 pairwise comparison as indicated (one-way ANOVA with Tukey’s post hoc test). **(D)** Hippocampal lysates were immunoprecipitated using an anti-PSD-95 antibody and Western Blots probed with a ZDHHC2 antibody. The ZDHHC2 input was first normalized to β-actin as a loading control. The amount of immunoprecipitated ZDHHC2 protein was then normalized to the β-actin normalized input. **(E)** 14 DIV, cultures were stimulated using cLTP treatment and PSD-95 palmitoylation determined using the Acyl-Rac assay. *P<0.05 (unpaired student’s *t*-test). **(F)** HA-ZDHHC2-8xA phospho-dead mutation reduces co-immunoprecipitation (co-IP) of PSD-95. The PSD-95 input was first normalized to β-actin as a loading control. The amount of immunoprecipitated PSD-95 protein was then normalized to the β-actin normalized input. **(G)** Hippocampal cells were either left untransfected or transfected with HA-ZDHHC2-WT or HA-ZDHHC2-8xA at time of plating. At 14 DIV endogenous PSD-95 palmitoylation was determined using the Acyl-Rac assay. ns = not significant. ***P < 0.001 vs first condition in bar chart. ##P < 0.01 pairwise comparison as indicated (one-way ANOVA with Tukey’s post hoc test). Results are mean ± s.e.m. with individual data points shown. N = 3-5 independent hippocampal cultures per experiment. **(H)** Schematic of activity-dependent changes in ZDHHC2 C-terminal phospho-regulation of PSD-95 substrate interactions.

Synaptic activity can alter ZDHHC2-mediated palmitoylation of its downstream substrate, PSD-95 (Fukata et al., 2013). We observed a decrease in the association between ZDHHC2 and PSD-95 1 hr after cLTP (Fig. 5D) which coincided with an increase in the palmitoylation of PSD-95 (Fig 5E). While appearing initially contradictory, this could reflect changes in the kinetics of ZDHHC2 or changes in the binding domain of ZDHHC2 and/or PSD-95 after activity-induced ZDHHC2 dephosphorylation. To determine whether activity-induced changes in ZDHHC2/PSD-95 interactions and PSD-95 palmitoylation are directly due to changes in ZDHHC2 phosphorylation, we transfected cells with either ZDHHC2 phospho-mimetic (8 Ser/Thr residues between 330-366 aa changed to Asp/Glu; HA-ZDHHC2-8xDE) or phospho-dead (8 serine/threonine residues between 330-366 aa changed to alanines; HA-ZDHHC2-8xA; Salaun et al., 2017) constructs. These C-terminal serine and threonine residues were specifically targeted as phosphorylation of these sites have been shown to be important for membrane localization (Salaun et al., 2017) and as synaptic activity can alter the membrane localization of ZDHHC2 (Fukata et al., 2013). In line with our observed decrease in ZDHHC2 phosphorylation and ZDHHC2/PSD-95 interaction following cLTP, there was a robust decrease in the association of PSD-95 with the HA-ZDHHC2-8xA phospho-dead mutant compared to HA-ZDHHC2-WT (Figure 5F). The phospho-mimetic HA-ZDHHC2-8xDE mutant did not increase ZDHHC2/PSD-95 association when compared with HA-ZDHHC2-WT, suggesting high basal phosphorylation of these Ser and Thr residues (Figure 5F). To determine whether decreased ZDHHC2 phosphorylation drives increased PSD-95 palmitoylation, we overexpressed HA-ZDHHC2-WT or the phospho-dead HA-ZDHHC2-8xA mutant. Both constructs increased the amount of palmitoylated PSD-95 equally (Figure 5G), possibly because overexpression of either construct is sufficient to achieve saturated PSD-95 palmitoylation.

Overall, these results indicate that cLTP decreases the phosphorylation of ZDHHC2 at its C-terminal tail, resulting in a decrease in the association between PSD-95 and ZDHHC2 and an increase in PSD-95 palmitoylation (Figure 5H).

### *In vivo* regulation of ZDHHCs following fear conditioning

Having identified a number of cLTP-induced post-translational modifications for ZDHHC enzymes *in vitro*, we next tested whether similar post-translational modifications occurred *in vivo* in response to a hippocampal-dependent learning event. Hippocampal lysates were collected 1 hr after contextual fear conditioning on adult male mice and changes in total protein, phosphorylation and palmitoylation were assayed (Figure 6). In line with our *in vitro* results, we observed a significant decrease in ZDHHC5 protein levels after fear conditioning (Input, Fig 6A), in line with the activity-dependent decrease in ZDHHC5 protein stability observed after cLTP *in vitro* (Fig. 2A, B). However, this decrease was not accompanied by a relative increase in phosphorylation as seen *in vitro* (Fig 6A) or palmitoylation (Fig 6B), perhaps due to differences in cell type composition of the two samples, or stimulation paradigms. No significant changes were observed for ZDHHC8 in any measure following FC (Fig. 6C, D), in line with *in vitro* findings (Fig. 3). We did not observe changes in ZDHHC9 palmitoylation following FC, indicating that the activity-dependent decrease in ZDHHC9 palmitoylation may be specific to certain types of synaptic stimuli (Fig. 6E). Finally, at 40 min post cLTP *in vitro*, we observed a significant reduction in ZDHHC2 phosphorylation (Figure 6G), along with no changes in ZDHHC2 palmitoylation or total protein levels (Figure 6G, H). These observations indicate that several of the post-translational changes observed with cLTP *in vitro* are mirrored following a learning event *in vivo*.

**Figure 6.**
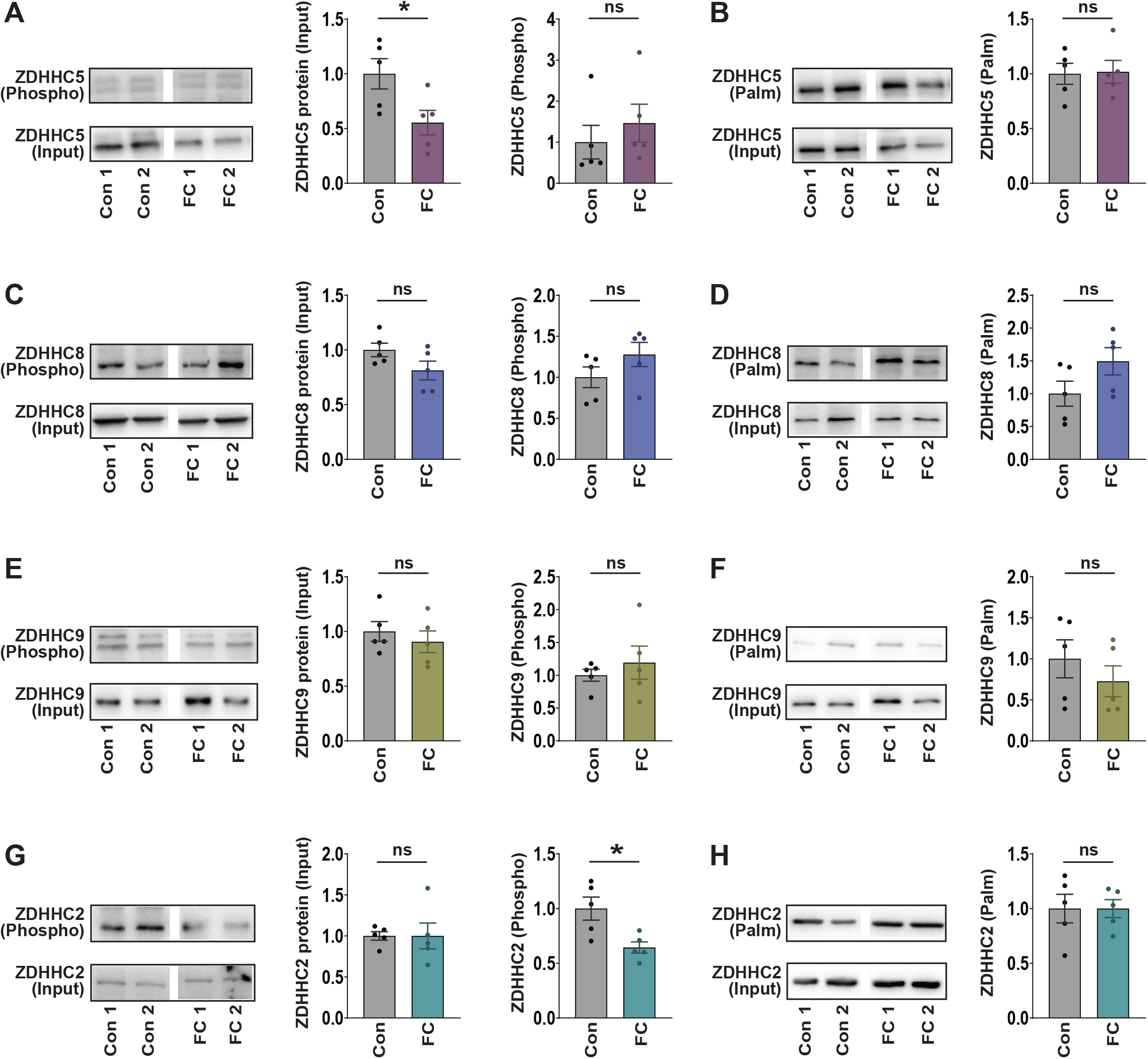
ZDHHC activity-dependent post-translational changes *in vivo*. 9-week-old male mice were subjected to contextual fear conditioning (cFC) and hippocampal lysates from conditioned and unconditioned mice collected 1 hour later. **(A, C, E, G)** Lysates were run through the phospho-protein purification assay and Western Blots probed with ZDHHC5, ZDHHC8, ZDHHC9 and ZDHHC2, respectively. Input fractions demonstrate overall ZDHHC protein levels in control and cFC lysates **(B, D, F, H)** Acyl-Rac assay showing palmitoylated ZDHHC5, ZDHHC8, ZDHHC9 and ZDHHC2, respectively, in control or cFC treated mice. ns = not significant, *P<0.05 (unpaired student’s *t*-test). Results are mean ± s.e.m. with individual data points shown. N = 5 hippocampi per experiment.

### cLTP does not alter APT2 or ABHD17 transcription, protein turnover, PTMs or activity

Activity-dependent changes in synaptic substrate palmitoylation might also be a result of dynamic regulation of depalmitoylating enzymes. To directly investigate whether synaptic activity impacts depalmitoylating enzyme activity, we utilized a recently developed probe (depalmitoylation probe-5 or DPP-5) that generates fluorescent signal in response to thioestaerase activity in live cells (Qiu et al., 2018). We first validated that the DPP-5 probe is sensitive enough to detect changes in thioesterase activity in neurons using the pan-thioesterase inhibitor palmostatin B (Suppl. Fig. 2), and then tested whether thioesterase activity is altered following cLTP induction. We observed no significant changes in DPP-5 fluorescence either 1 hr (Fig. 7A, C, D) and 24 hrs (Fig. 7B, E, F) after cLTP treatment, indicating that there is no change in thioestaerase activity following cLTP. As the membrane-anchored serine hydrolases, ABHD17A, 17B, and 17C, have the highest de-palmitoylating activity against PSD-95 in neurons (Yokoi et al., 2016), we focused our assays on these enzymes. We observed no activity-induced changes in ABHD17 total protein (phospho assay input, P = 0.858, Fig. 7G), palmitoylation (Fig. 7G) or phosphorylation (Fig. 7H) (ABHD17 antibody validation in Suppl Fig 1). Together, these data suggest that activity-dependent changes in PSD-95 palmitoylation are mediated through post-translational regulation of ZDHHC2 as opposed to regulation of ABHD17 function. We also assessed post-translational modifications of APT2, which is expressed in hippocampal neurons (Wild et al., 2022) and again found no changes in APT2 protein levels (Fig 7J; phospo-assay input, P = 0.753), palmitoylation (Fig. 7I), or phosphorylation (Fig 7J). In summary, we did not find evidence of cLTP induced activity-dependent changes in de-palmitoylating enzyme activity or post-translational regulation, indicating that the ZDHHC enzymes may be the primary regulators of synaptic activity-induced differential palmitoylation of substrate proteins.

**Figure 7.**
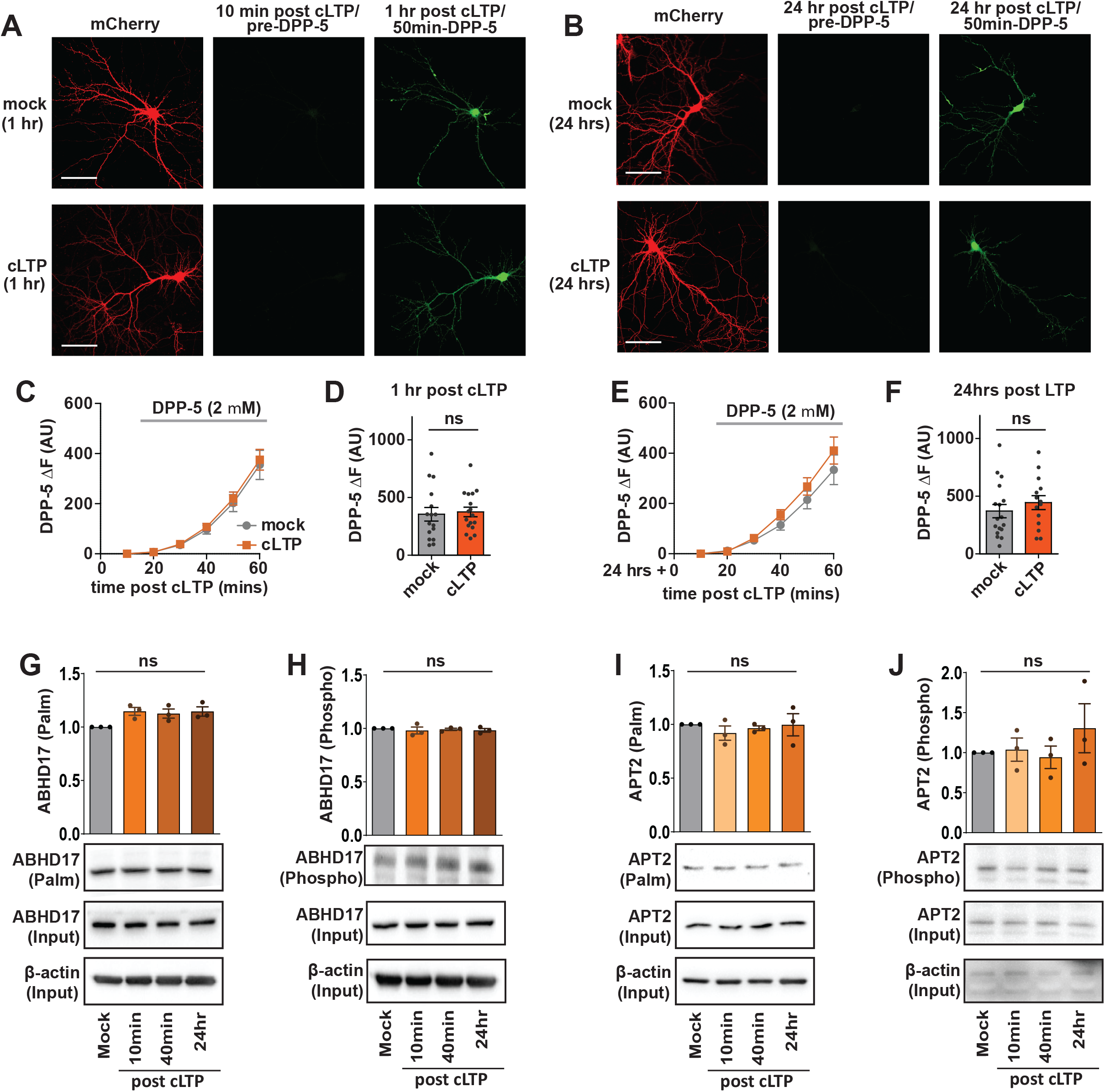
De-palmitoylating enzyme activity unchanged following cLTP. **(A)** Representative images of fluorescent depalmitoylation probe-5 (DPP-5; 2 μM) 1 hour after cLTP treatment in 14 DIV cultured hippocampal neurons transfected at 11 DIV with mCherry. *Left*: mCherry cell fill. *Middle*: Background fluorescence within mCherry mask prior to addition of DPP-5 to the bath. *Right*: DPP-5 fluorescence within mCherry mask 50 mins post addition of DPP-5 to the bath (1 hr post cLTP). Scale bar = 100 μm. **(B)** As (A) but 24 hrs post cLTP treatment. **(C)** Graph of time-course of DPP-5 fluorescence (ΔF) increase following cLTP and subsequent bath addition of DPP-5. No significant difference was observed between mock and cLTP treated neurons at any time-point. **(D)** Graph of DPP-5 fluorescence (ΔF) 1 hr post cLTP and 50 min post addition of DPP-5 to the bath. No significant difference was observed between mock and cLTP treated neurons at any time-point. **(E, F)** As (C) and (D) but 24 hrs post cLTP and 50 min post addition of DPP-5 to the bath. ns = not significant (unpaired student’s *t*-test). Results are mean ± s.e.m. with individual data points shown. N = 14-17 neurons from 2-4 hippocampal cultures. **(G)** Acyl-Rac assay showing palmitoylated ABHD17 in cultured hippocampal neurons (14 DIV) following cLTP. **(H)** Phospho-protein purification assay showing phosphorylated ABHD17 following cLTP. **(I)** Acyl-Rac assay showing palmitoylated APT2 in cultured hippocampal neurons (14 DIV) following cLTP. **(J)** Phospho-protein purification assay showing phosphorylated APT2 following cLTP. ns = not significant (one-way ANOVA with Tukey’s post hoc test). Results are mean ± s.e.m. with individual data points shown. N = 3 hippocampal cultures per experiment.

## Discussion

While accumulating evidence indicates that the dynamic palmitoylation of many synaptic proteins occurs in response to changes in synaptic activity and is critical for synaptic plasticity, the mechanism by which this occurs is largely unknown. Given the reversible nature of palmitoylation, changes in protein palmitoylation might reflect changes in palmitoylation and/or depalmitoylation rates. Our study set out to elucidate the post-translational modifications regulating the palmitoylation machinery, that might ultimately modulate enzyme function and activity-induced palmitoylation of synaptic substrates. We performed a systematic screen of endogenous post-translational modifications for numerous neuronal palmitoylating and depalmitoylating enzymes, as well as an investigation of how these modifications might alter protein function or substrate interactions.

### cLTP induces changes in post-translational modifications of ZDHHC5, but not ZDHHC8

Our previous work demonstrated that glycine cLTP induction results in the transient removal of ZDHHC5 from the synaptic membrane, followed by a 2-fold increase in ZDHHC5 membrane localization 20 mins later (Brigidi et al., 2015). These changes in trafficking were accompanied by increased δ-catenin palmitoylation and AMPAR stabilization at the synapse. Here, we also demonstrate a 2.5-fold decrease in total ZDHHC5 protein at the same time point that is a result of activity-dependent phosphorylation of Ser and Thr residues (Ser-569, Ser-572 and Thr-573) within the polo-box domain of the ZDHHC5 C-terminal. How then might both increased synaptic localization, and overall protein degradation of ZDHHC5 work cooperatively to alter synaptic function? Recruiting ZDHHC5 to the synaptic membrane and degrading non-synaptic ZDHHC5 could profoundly impact the palmitoylation of its downstream substrates that are localized to different compartments. For example, while palmitoylation of δ-catenin stabilizes AMPAR at the synaptic membrane, palmitoylation of neuronal scaffolding protein GRIP1B by ZDHHC5 enhances AMPAR turnover (Thomas et al., 2012). It is therefore interesting to speculate that enhanced synaptic activity might differentially regulate ZDHHC5 to stabilize AMPARs by simultaneously increasing the palmitoylation of δ-catenin and decreasing the palmitoylation of GRIP1B in different subcellular compartments.

We also observed an increase in ZDHHC5 palmitoylation following cLTP. Several Cys residues in the C-terminal of ZDHHC5 are known to be palmitoylated (Cys-236, -237 and -245; human isoform), that lie close to a juxtamembrane amphipathic helix. This helix forms part of the binding site for ZDHHC5 accessory protein GOLGA7B (Woodley and Collins, 2019) and the Na-K ATPase Na-pump (Plain et al., 2020). Furthermore, ZDHHC20 was identified as the enzyme that can palmitoylate these residues in ZDHHC5 in HEK 293 cells (Plain et al., 2020). It is therefore possible that activity-dependent palmitoylation of ZDHHC5 could be mediated by ZDHHC20, or alternatively by ZDHHC enzymes that co-localize with ZDHHC5 in neurons that are yet to be identified. Previous work from our group found that mutation of the aforementioned C-terminal ZDHHC5 Cys residues increased surface localization of ZDHHC5 in rat hippocampal neurons (Shimell et al., 2019), raising the possibility that activity-dependent increases in ZDHHC5 palmitoylation could be another means to facilitate ZDHHC5 endocytosis and re-localization.

Surprisingly, we did not observe any activity-dependent post-translational modifications for ZDHHC8, an enzyme which shares a high degree of sequence homology with ZDHHC5 (60 %), and which might therefore be expected to be responsive to neuronal activity. ZDHHC5 contains many regulatory motifs that are also found in ZDHHC8, including a C-terminal PDZ binding domain (EISV; Thomas et al., 2012), tyrosine endocytic motif (YDNL; Brigidi et al., 2015) and a Polo-box like domain (ZDHHC8 sequence: DSGVYDT). Additionally, the two enzymes are functionally redundant in palmitoylating common substrates such as ankyrin-G in polarized epithelial cells (ANK3; He et al., 2014). However, ZDHHC5 and ZDHHC8 show distinct dendritic sub-cellular localizations in neurons (Thomas et al., 2012), and analysis of *Zdhhc5* and *Zdhhc8* expression in the mouse brain revealed that they are differentially enriched within regional neuronal sub-populations of the hippocampus and cortex (Wild et al., 2021). Our findings here that ZDHHC5 and ZDHHC8 are differentially responsive to neuronal activity in the hippocampus support the notion that these two enzymes may also have distinct roles in regulating neuronal function and synaptic plasticity.

Previous studies have reported activity-dependent phospho-regulation of ZDHHC5 and ZDHHC8, including a recent study which revealed that phosphorylation at Tyr-91 within the intracellular catalytic loop by LYN kinase in adipocytes reduces ZDHHC5 enzymatic activity (Hao et al., 2020). However, it is not yet clear if this modification occurs in neurons, or if it is dynamically regulated by neuronal activity. A recent phospho-proteomic screen identified several Ser residues in rat ZDHHC5 that were differentially phosphorylated following changes in neuronal activity, including Ser-380, Ser-432 and Ser-621, as well as 10 other Ser/Thr phophosites that were not altered by activity (Desch et al., 2021). Interestingly, aside from ZDHHC5, the only other ZDHHC that was identified to be differentially phosphorylated following activity changes in neurons was at a single residue in ZDHHC8 (Ser-335; Desch et al., 2021). A previous study also found that BDNF/TrkB signaling can stimulate PKMζ phosphorylation of ZDHHC8 in cortical neurons, indicating that activity-dependent phospho-regulation of ZDHHC8 is possible during different types of neuronal activity (Yoshii et al., 2011). Overall, we have found in this study that numerous activity-dependent post-translational modifications regulate ZDHHC5, supporting previous studies that have identified ZDHHC5 as a key mediator of dynamic synaptic substrate palmitoylation following changes in neuronal activity.

### ZDHHC9 palmitoylation is decreased following cLTP

We found that cLTP in can induce a decrease in palmitoylation within the ZDHHC9 catalytic DHHC domain. This is accompanied by a decrease in the palmitoylation of known ZDHHC9 substrates TC10 and NRAS (Shimell et al., 2019), indicating a potential role for activity-regulated control of ZDHHC9 enzymatic activity. Previous studies from our group revealed that knock-down or knock-out of ZDHHC9 reduces the palmitoylation of TC10 and NRAS (Shimell et al., 2019), supporting the notion that decreased substrate palmitoylation may be a direct result of reduced ZDHHC9 function. An important outstanding question is how might neuronal activity lead to decreased ZDHHC9 palmitoylation? One potential mechanism is altered activity of ZDHHC9 palmitoylating or depalmitoylating enzymes, that might in turn alter ZDHHC9 palmitoylation. Alternatively, because the decreased palmitoylation appears to occur within the catalytic domain of the protein, it is possible that neuronal activity might alter the interaction between ZDHHC9 and its accessory protein GOLGA7, which is known to stabilize palmitoylation of ZDHHC9 within the catalytic ‘DHHC’ domain (Mitchell et al., 2014; Swarthout et al., 2005). Other mechanisms that alter ZDHHC9 autopalmitoylation within the active site may also be responsible. Our observation of activity-dependent regulation of ZDHHC9 is particularly interesting, given the important role that this enzyme plays in neuronal outgrowth and synapse formation, two processes that are impaired when the catalytic Cys residue of ZDHHC9 is mutated to Ser (Shimell et al., 2019). Future work is needed to determine the functional consequences of activity-dependent regulation of ZDHHC9 palmitoylation.

### ZDHHC2 phospho-regulation alters substrate interactions

In this study, we observed an increase in PSD-95 palmitoylation following cLTP as previously reported (Brigidi et al., 2015; Nasseri et al., 2021). However, we also observed a reduction in ZDHHC2 / PSD-95 interactions following cLTP which may initially appear counter-intuitive. One explanation is that changes in the phosphorylation of ZDHHC2 following cLTP impacts ZDHHC2 enzyme kinetics, resulting in a more rapid transfer of palmitic acid to substrate proteins and hence a decrease in the duration of interaction with substrates. It is also possible that changes in ZDHHC2 phosphorylation changes the site(s) of ZDHHC2/PSD-95 interaction, resulting in less interaction but greater palmitoylation.

### *In vivo* changes in ZDHHC post-translational modifications

Many of the ZDHHC post translational changes we observed following cLTP *in vitro* were also replicated following cFC *in vivo*, including a striking decrease in both ZDHHC5 total protein and ZDHHC2 phosphorylation. Recent work has revealed that relatively few neurons and synapses in the hippocampus undergo consolidative plasticity to become so called ‘engram’ neurons following learning stimuli such as cFC (Denny et al., 2014; Josselyn and Tonegawa, 2020; Liu et al., 2012). However, it has also been reported that in addition to cell autonomous mechanisms regulating engram formation, numerous non-engram cells are activated following cFC, including non-engram excitatory neurons, inhibitory neurons, neural progenitors and glia (Denny et al., 2014; Li et al., 2020; Pan et al., 2020; Seo et al., 2015; Stefanelli et al., 2016). Given the magnitude of the changes we have observed here, our results would indicate that ZDHHC post-translational changes extend beyond the relatively small population of engram neurons, to the wider network of hippocampal cells activated following cFC that support engram formation. Furthermore, our findings support the notion that multiple types of synaptic stimuli can drive similar changes in ZDHHC post-translational modifications.

### Evidence for activity-dependent regulation of palmitoylating, but not depalmitoylating enzymes

Despite reports of both activity-dependent increases and decreases in palmitoylation in the hippocampus (Nasseri et al., 2021), evidence of activity-dependent regulation of the family of depalmitoylating enzymes is currently lacking. The findings in this study are consistent with a recent phospho-proteomic screen that did not report activity-dependent changes in the phosphorylation of any of the best characterized depalmitoylating enzymes, including APT1, APT2, PPT1 and all members of the ABHD family (Desch et al., 2021). It is possible that more detailed future studies might reveal novel mechanisms that regulate the family of depalmitoylating enzymes and promote activity-dependent changes in substrate palmitoylation. However, given the current evidence, we propose that the ZDHHC family of proteins are the primary sensors of neuronal activity, and that their bi-directional regulation is the effector of both increases and decreases in neuronal substrate palmitoylation.

## Methods

### DNA constructs and primers

N-terminal HA-tagged mouse DHHC 1-24, HA-tagged ZDHHC5 AAA, HA-tagged P35, and Myc-tagged CDK5 plasmids were kind gifts from Dr. Gareth M. Thomas (Temple university, Philadelphia, Pennsylvania). FLAG-tagged-ABHD17A, 17B, and 17C were kind gifts from Dr. Elizabeth Conibear (University of British Columbia, Vancouver, BC). Myc-tagged PLK2, and Myc-tagged PLK2 kinase-dead mutant were kind gifts from Dr. Daniel Pak (Georgetown University, Washington, DC). shRNA against ZDHHC5 was a kind gift from Dr. Richard Huganir (Johns Hopkins University, Baltimore, MD).

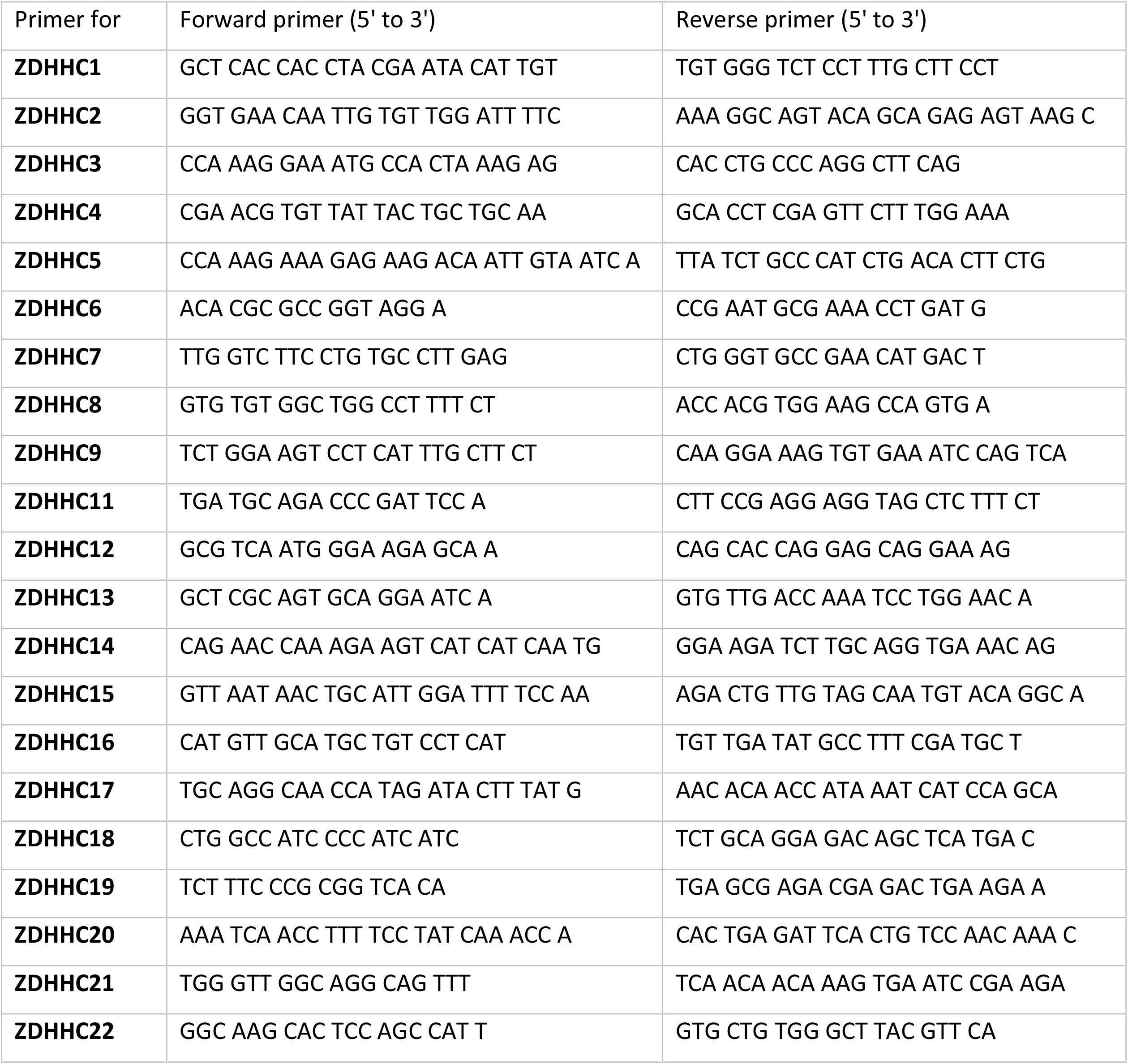
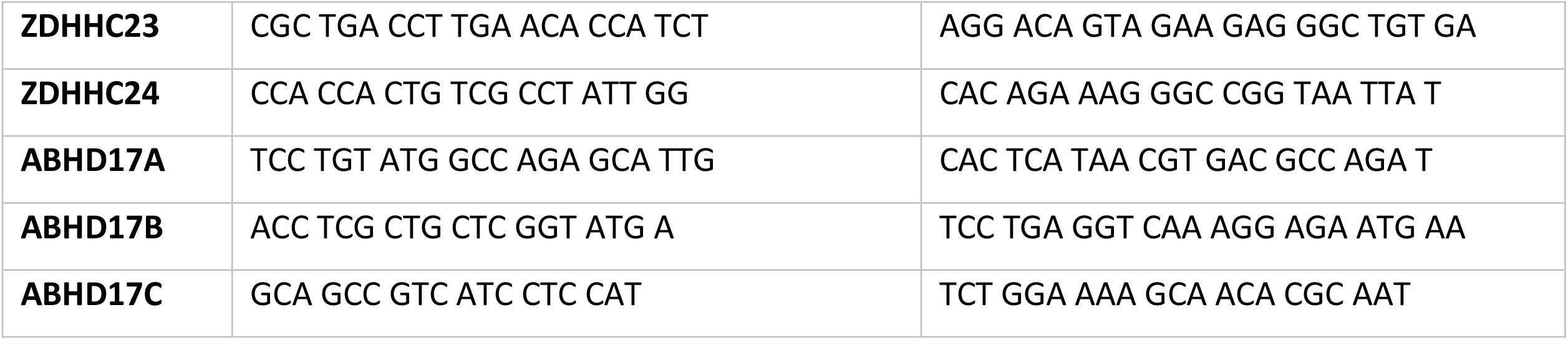

### Antibodies

#### Primary antibodies used

β-actin (1:5000, Sigma A1978), anti ZDHHC1 (1:1000, Abcam ab223042), anti ZDHHC2 (1:1000, Santa Cruz Biotechnology sc-515204), anti ZDHHC2 (1:500, Sigma SAB1101457), anti ZDHHC3 (1:500, Aviva Systems Biology ARP59576), anti ZDHHC3 (1:500, Sigma SAB2107413), anti ZDHHC3 (1:1000, Abcam ab124084),anti ZDHHC3 (1:1000, Abcam ab31837), anti ZDHHC4 (1:500, Aviva Systems Biology ARP78440), anti ZDHHC5 (1:1000 for WB, 5µg for IP, Sigma HPA014670), anti ZDHHC6 (1:600, Abcam ab121423), anti ZDHHC7 (1:500, Aviva Systems Biology OAAB11570), anti ZDHHC7 (1:500, BosterBio A11785), anti ZDHHC8 (1:500, Santa Cruz Biotechnology sc-374191), anti ZDHHC9 (1:1000, Sigma SAB4502104), anti ZDHHC9 (1:1000, Thermo Fisher Scientific PA5-26721), anti ZDHHC11 (1:500, Abcam ab116065), anti ZDHHC12 (1:500, Aviva Systems Biology ARP60674), anti ZDHHC13 (1:500, Aviva Systems Biology ARP44398), anti ZDHHC14 (1:500, Aviva Systems Biology ARP42628), anti ZDHHC15 (1:500, Sigma SAB4500608), anti ZDHHC15 (1:200, Abcam ab121203), anti ZDHHC15 (1:500, Santa Cruz

Biotechnology sc-169847), anti ZDHHC15 (1:500, Thermo Fisher Scientific PA5-39327), anti ZDHHC16 (1:500, Aviva Systems Biology ARP50063), anti ZDHHC17 (1:300, Proteintech 15465-1-AP), anti ZDHHC17 (1:500, Sigma AV47141), anti ZDHHC18 (1:1000, Abcam ab154790), anti ZDHHC19 (1:500, Abcam ab179545), anti ZDHHC20 (1:500, Aviva Systems Biology ARP72069), anti ZDHHC21 (1:300, Abcam ab103755), anti ZDHHC22 (1:500, Santa Cruz Biotechnology sc-514005), Phospho-PLK Binding Motif (ST*P) (1:1000, Cell Signaling Technology 5243S), anti-HA (1:1000, Cell Signaling Technology C29F4), anti-myc (1:1000, Cell Signaling Technology 2276), anti-GFP (1:3000, Abcam ab290), anti ABHD17 (1:1000, Origene TA331704), anti ABHD17 (1:1000, Proteintech 15854-1-AP), anti-FLAG (1:1000, Sigma F7425), PSD-95 (1:500, Abcam, ab2723), APT2 (1:500, Abcam, ab151578), TC10 (1:1000, Abcam, ab168645), N-Ras (1:500, Santa Cruz Biotechnology sc-31).

#### Secondary antibodies used

Goat anti-mouse IgG-HRP (1:6000, BioRad 170-6516), Goat anti-rabbit IgG-HRP (1:6000, BioRad 170-6515)

### Cell culture

#### Primary hippocampal neurons

Hippocampi from embryonic day 18 (E18) Sprague Dawley rats of either sex were prepared as previously described (Xie et al., 2000). Briefly, hippocampi were dissected, and incubated with 0.25% Trypsin (Thermo Fisher Scientific) and 0.05% DNase (Thermo Fisher Scientific) for 20 and 3 minutes, respectively. Cells were dissociated with titration and plated at a density of 3.2 million/10-cm culture dish for biochemical assays. Cells were allowed to adhere in plating media containing Minimum Essential Media (MEM; Gibco, Thermo Fisher Scientific), supplemented with 10% (vol/vol) heat-inactivated-fetal bovine serum (FBS) (Gibco, Thermo Fisher Scientific), sodium pyruvate (Gibco, Thermo Fisher Scientific), 0.5% glucose, GlutaMAX (Gibco, Thermo Fisher Scientific), and Pen/Strep (Gibco, Thermo Fisher Scientific). After 3 hours plating media was replaced with maintenance media containing Neurobasal medium (Gibco, Thermo Fisher Scientific), supplemented with NeuroCult SM1 (StemCell, instead of B27 in the original protocol), GlutaMAX (Gibco, Thermo Fisher Scientific), and Pen/Strep (Gibco, Thermo Fisher Scientific). Cultures were maintained at 37°C and 5% CO2.

#### HEK Cells

HEK293T cells (Sigma) were aliquoted into a 10 cm culture dish with 15 ml of pre-warmed (37°C) DMEM (Gibco, Thermo Fisher Scientific), supplemented with 10% (vol/vol) heat-inactivated fetal bovine serum (FBS) (Gibco, Thermo Fisher Scientific), 1% Pen/Strep. HEK Cells were maintained in an incubator at 37°C and 5% CO2.

### Transfection

#### Primary hippocampal cultures - transient transfections

Neurons were transfected at 9-11 days in vitro (DIV) using Lipofectamine 2000 (Invitrogen) according to the manufacturer’s protocol and used for experiments on DIV 12-15.

#### Primary hippocampal cultures – Amaxa nucleofection

Neurons were nucleofected with identified plasmids prior to plating at 0 DIV using Amaxa Rat Neuron Nucleofector kit (DGP-1003; Lonza) according to manufacturer’s protocol. Cells were then used for experiment at 13-15 DIV.

#### HEK Cells

HEK293T Cells were transfected at 70-80% confluency, using Lipofectamine 2000 (Invitrogen) according to the manufacturer’s recommendations and used for experiments 24-48 hours after transfection.

### Neuronal stimulation (cLTP)

Neuronal activity was enhanced as per previously published protocol (Lu et al., 2001). Briefly, at 13-15 days *in vitro*, the maintenance media was removed and stored at 37°C and cells washed 3 times with pre-warmed (37°C) Mg^2+^-free extracellular solution made of 140 mM NaCl, 1.3 mM CaCl_2_, 5.0 mM KCl, 25 mM HEPES, and 33 mM glucose, supplemented with 0.0005 mM TTX and 0.001 mM strychnine (pH 7.4). To chemically induce LTP, cells were incubated with the above extracellular solution supplemented with 200 µM glycine for 3 minutes. Cells were then washed 2 times with the above extracellular solution containing 2 mM MgCl_2_, and then replaced with stored maintenance media. Neuronal cells were maintained in 37°C incubator with 5% CO_2_ for the indicated time prior to experimentation. Control cells were treated the same as experimental groups but were not exposed to glycine during the 3-minute incubation.

### RNA isolation and qPCR

At 15 DIV hippocampal cultured neurons were stimulated as described above and mRNA was isolated after identified time points using TRIzol Reagent (ThermoFisher Scientific) as described by manufacturer’s instructions. 200 ng of total DNA-free RNA was reverse transcribed using Verso cDNA Synthesis Kit (Thermo Scientific). The cDNA was then quantified by qPCR using SYBR green (ThermoFisher Scientific). Real-time quantitative PCR (qPCR) analysis was performed at the Biomedical Research Center at UBC using a 7900HT Real-Time PCR thermocycler machine (Applied Biosystems). mRNA levels of genes of interest were normalized to GAPDH and shown as fold change over baseline using the delta-delta CT method (Schmittgen and Livak, 2008).

### Western blot assay

Brain tissue, primary hippocampal neurons, and HEK293T cells were washed with ice-cold PBS and lysed in ice-cold Tris Lysis Buffer containing 1% IGEPAL (Sigma), 50mM Tris-HCl pH 7.5, 150mM NaCl, 10% Glycerol, supplemented with phenylmethanesulfonyl fluoride solution (PMSF) and a protease inhibitor cocktail with EDTA (Roche). The samples were vortexed, and run through a 26-gauge syringe and kept at 4°C to nutate for 30 minutes. Lysates were then cleared by spinning down at 16,000 x g for 30 minutes at 4°C. Protein quantification was done using a BCA assay kit (Thermo Scientific) as per the manufacturer’s instructions. Proteins were separated by electrophoresis on a 10-12% SDS-PAGE gel. Proteins were then transferred to a PVDF membrane (BioRad) and blocked for one hour in 3-5% BSA in TBST. The membrane was then incubated overnight at 4°C with identified primary antibody. The membranes were then washed times for 15 minutes in TBST at room temperature with agitation and incubated with appropriate secondary antibodies for 1 hour at room temperature, before being washed 3 times for 15 minutes with TBST. Proteins were visualized using chemiluminescence (Immobilon Western Chemiluminescent HRP Substrate, Millipore, #WBKLS0500) on Bio-Rad ChemiDoc (XRS**+**). Blots were quantified using Image J software. For reprobing, blots were stripped as per previously published protocol (Yeung and Stanley, 2009). For western blot analysis, the protein of interest input was first normalized to β-actin as a loading control. For Acyl-Rac palmitoylation and phospho-protein assays, the amount of palmitoylated or phospho-protein was then normalized to the β-actin normalized input.

### Immunoprecipitation

Cells were lysed as described above and incubated overnight at 4°C with antibodies under gentle rotation. 80-100 μL of a mix of protein A and G-Agarose (Roche) was added to the samples, beads were recovered 4 hrs later and then washed 5 times with cold Tris Lysis Buffer. Proteins were eluted from beads by heating in 2X SDS loading buffer for 5 min at 80°C. Samples were analyzed by SDS-PAGE, then immunoblotted with identified antibodies.

### Acyl-Rac Assay

Protein palmitoylation assay was performed using CAPTUREome S-palmitoylated protein kit (Badrilla, Leeds, UK), as described by manufacturer’s protocol, with the following modification: Protein concentration was measured after dissolving the precipitated protein, to ensure starting with equal protein concentrations. Hippocampal tissue, or cultured neurons were lysed and incubated with blocking reagent to block all free thiol groups. Extracted proteins were then acetone precipitated. Pellets were re-dissolved and protein concentration was measured using BCA assay. The palmitate groups on proteins were cleaved using thioester cleavage reagent. Proteins with newly liberated thiols were then captured using CAPTUREome resin. Captured proteins were then eluted from resin. samples were analyzed by SDS-PAGE, then immunoblotted with identified antibodies.

### Phospho-Protein Purification Assay

Protein phosphorylation assay was performed using PhosphoProtein Purification Kit (Qiagen), exactly as described by manufacture’s guideline. For negative control, the lysates were incubated with 800 units of lambda protein phosphatase (New England Biologicals) for 45 minutes at room temperature.

### Context-dependent fear conditioning

9-week-old male mice were first habituated by handling 15 min per day for 3 days. On the training day, mice were placed in conditioning chamber designed by CleverSys (CleverSys Inc.) with a shock floor and habituated for 2 min. The conditioned group then received a 0.3 mA foot shock for 5 s, whereas the control group did not. 1 hr later, mice were placed back into the same conditioning chamber for 5 min, and the total freezing time quantified using FreezeScan software by CleverSys Inc. Immediately after testing mice were euthanized and hippocampi isolated and used in Acyl-Rac and phospho-protein purification assays for further palmitoylation and phosphorylation analysis.

### Live-cell imaging of thioesterase activity with DPP-5

Cultured hippocampal neurons were transfected on 12-13 DIV with mCherry (0.8 μg). For experiments with Palmostatin B (PalmB), either control (DMSO 1/1000) or PalmB (5 μM; from 5mM stock in DMSO) solutions were added to the culture media 30mins prior to the experiment and were included throughout the imaging experiment. Mock or cLTP treatment was performed as described above. At the timepoints specified following treatment, neurons were transferred to an imaging chamber at 20°C with an artificial CSF (aCSF) media containing the following in mM: 135 NaCl, 5 KCl, 25 HEPES, 10 glucose, 2 CaCl2, 1 MgCl2, pH 7.4. Time-lapse images were acquired using a Zeiss LSM 880 AxioObserver Airyscan microscope with an 20X air objective using a 568 nm laser (mCherry) or a 488 nm laser (DPP-5) with AiryScan Fast mode. Two-color z-stack images of the soma and dendritic arbor covering a 422.0 × 422.0 μm field of view were acquired every 10 mins for a total of 50 mins. After the first acquisition in the time-lapse series to measure background green fluorescence, DPP-5 (2 μM) was added to the imaging chamber. Images were maximum intensity projected prior to analysis. To measure the amplitude of DPP-5 fluorescence changes, a binary mask was drawn of the soma and dendrites from the mCherry cell-fill that was then converted into a region of interest (ROI) to measure DPP-5 fluorescence changes. Data are reported as the change in DPP-5 fluorescence signal in the background subtracted mask (ΔF) in arbitrary units (AU).

### Statistical analysis

All data values are expressed as means ± SEM. Unless otherwise noted, statistical analysis was done using unpaired Student’s t-test and one-way ANOVA (with Dunnett’s multiple comparisons, or Tukey’s multiple comparisons) where applicable and defined when p < 0.05. In all figures, * = p < 0.05, ** = p < 0.01, *** = p < 0.001 and **** = p < 0.0001. All statistical analysis was performed in GraphPad Prism (La Jolla, CA, USA). Figures were generated using Adobe Illustrator CS6 software (Adobe Systems Inc., San Jose, CA).

## Acknowledgments/funding

This work was supported by a Canadian Health Services Research Foundation (F18-00650 CIHR Foundation Grant) to S.X.B, and by the National Institute of General Medical Sciences of the National Institutes of Health (R35 GM119840) to B.C.D.

---

**Supplementary Fig. 1.**
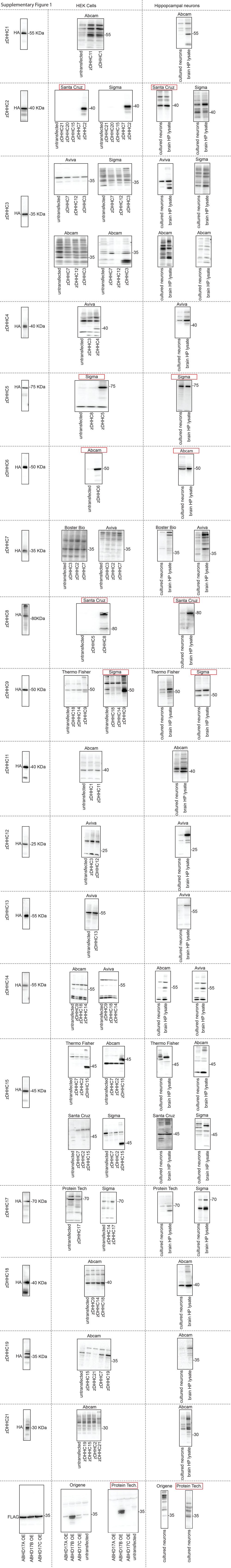
Validation of commercially available zDHHC antibodies. (A) Representative images of western blots testing efficacy and specificity of zDHHC and ABHD17 antibodies. Almost all commercially available antibodies for these enzymes were obtained, and the specificity of each antibody was tested. To do this, each zDHHC enzyme tagged with an HA epitope was transfected into HEK 293T cells. The closest phylogenetic or structural zDHHC for each enzyme was also transfected into HEK 293T cells in parallel. The efficacy and specificity of each antibody were then tested using western blotting. We also used Untransfected HEK cells as a negative control. We then tested the antibodies against endogenous proteins in lysates from either rat cultured hippocampal neurons or rat hippocampus Left: HEK cells expressing the indicated tagged zDHHC or ABHD and probed for the tag to demonstrate protein expression. Middle: HEK cells expressing the indicated tagged zDHHC and probed with an antibody from the indicated company. Right: Ability of the antibodies to detect endogenous zDHHCs or ABHDs in rat primary hippocampal cultures or hippocampal lysates. Transfection of zDHHC16, 20, and 23 was unsuccessful. Among all tested antibodies only five (marked with red boxes) were shown to be specific for the target proteins.

**Supplementary Fig. 2.**
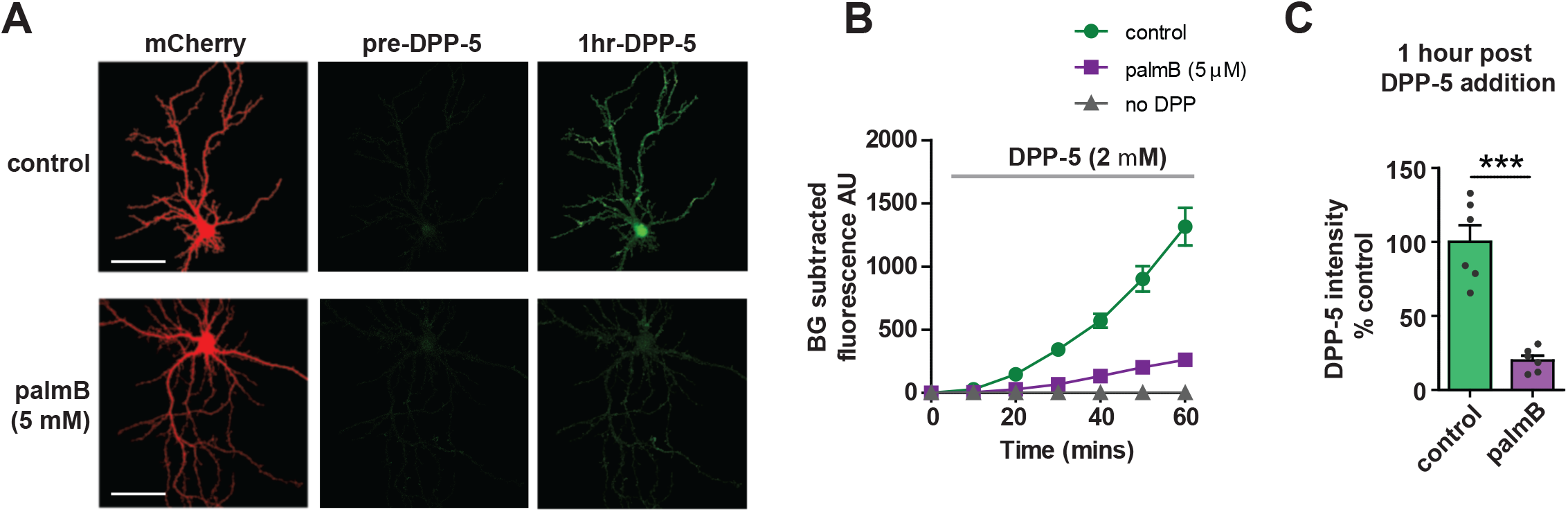
DPP-5 measures thioesterase activity in cultured hippocampal neurons. (A) Representative live-cell confocal images of DIV 14 cultured hippocampal neurons before and 1 hour after addition of DPP-5 (2 µM) +/−thioesterase inhibitor palmostatin B (PalmB, 5 µM). Left: mCherry cell fill. Middle: DPP-5 fluorescence in mCherry soma and dendrite mask pre-addition of DPP-5 (2 µM) to imaging chamber. Right: DPP-5 fluorescence in mask 1 hr post DPP-5 addition. Pan-thioesterase inhibitor PalmB substantially decreases DPP-5 fluorescence. Scale bar = 100 µm (B) Graph showing green flourescence within mCherry cell fill mask after addition of DPP-5 (2 µM) to the bath +/−PalmB. Triangles indicate background fluorescence timecourse without addition of DPP-5. (C) Graph showing percent inhibition of DPP-5 fluorescence by PalmB.

**Supplementary Fig. 3.**
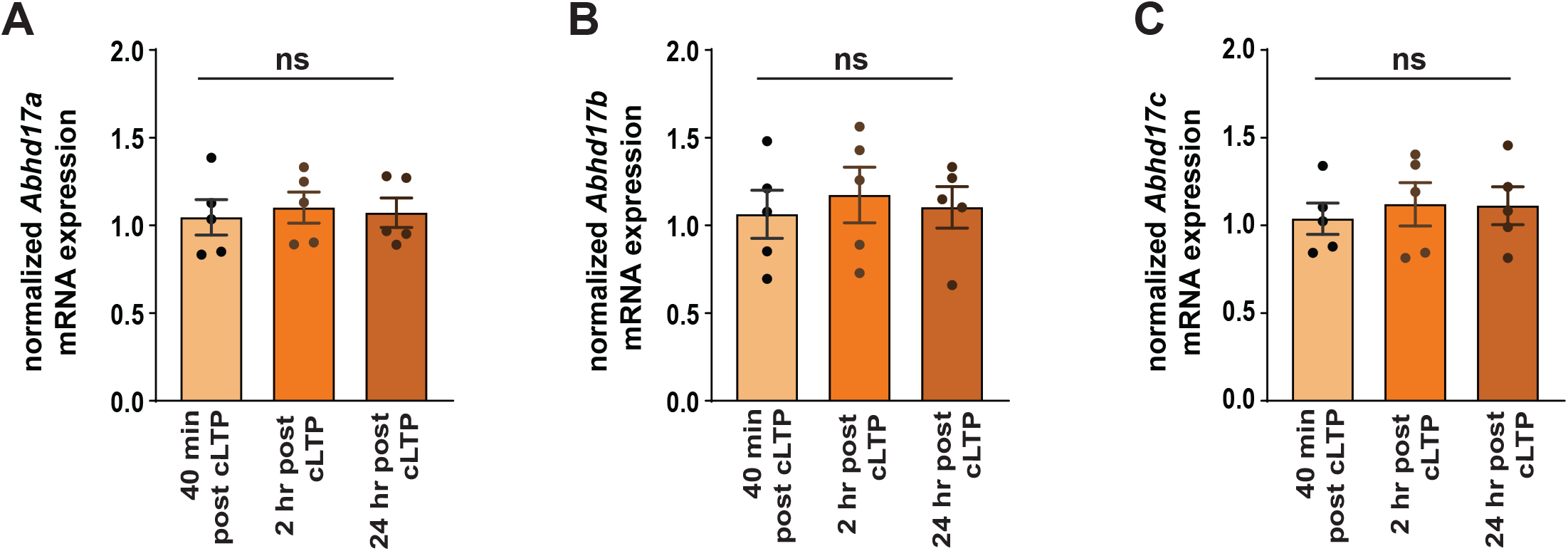
qRT-PCR of Abhd17a, b and c following cLTP. (A) Graph of qRT-PCR data from hippocampal culture lysates showing no change in *Abhd17a* mRNA expression following cLTP treatment at any timepoint. Data points are normalized to mock treated. (B) As (A) but for *Abhd17b*. (C) As (A) but for *Abhd17c*.

